# Cryo-electron microscopy revealed TACAN is extensively and specifically associated with membrane lipids

**DOI:** 10.1101/2021.10.11.463882

**Authors:** Zhen Wang, Fengying Fan, Lili Dong, Qingxia Wang, Yue Zhou, Rongchao Gao, Xuekui Yu

## Abstract

TACAN is not a mechanosensitive ion channel but significantly linked to the mechanical hyperalgesia. In this study, we show that the human TACAN is a homodimer with each monomer consisting of a body, a spring and a blade domains. The body domain contains six transmembrane helices that forms an independent channel. The spring domain adapts a loop-helix-loop configuration with the helix running within and parallel to the membrane. The blade domain is composed of two cytoplasmic helices. In addition, we found that all the helices of the body and the spring domains are specifically associated with membrane lipids. Particularly, a lipid core, residing within a cavity formed by the two body and spring domains, contacts with the helices from the body and spring domains and extends to reach two symmetrically arranged lipid clusters. These results extremely imply that the membrane lipids coordinate with the membrane-embedded protein to sense and transduce the mechanic signal.

## Introduction

A recent study has suggested that TACAN, also referred to as TMEM120A is a mechanosensory transduction ion channel and contributes to mechanosensitive currents in nociceptors and mechanical pain sensing in mice^1^. Another study has independently shown that attenuation of TACAN expression relieves the mechanical hyperalgesia produced by inflammatory mediators but not cancer-chemotherapeutic agents and implied that the TACAN could be a therapeutic target for pain induced by inflammatory conditions^2^. However, several groups have shown that TACAN is not a mechanosensitive ion channel, although they cannot ascertain its biological function^3,4,5^

The self-assembled lipid bilayer of plasma membrane possesses inherently large and anisotropic forces^6^. The changes of bilayer force would cause geometric changes of membrane-embedded proteins^6^. Lipid composition in cellular membrane are also known to influence the function of transmembrane proteins by modulation of membrane physicochemical properties and the lipid-protein interaction^7, 8^. Even several non-mechanically gated ion channels have been reported to be regulated by membrane stretches and lipid compositions^9, 10, 11, 12^. However, the mechanism how the change in membrane tension caused by the mechanical force is sensed and transduced remains elusive. Here, we show that the TACAN is extensively and specifically associated with membrane lipids, which provides a structural frame work to understand the mechanism of mechanical sensing.

## Results

### Overall structure of TACAN

We expressed and purified the full-length human TACAN (343 amino acids). The protein displayed good behavior as illustrated by gel filtration, SDS-PAGE and cryo-EM imaging (Supplementary Fig. S1a-c). Through image processing as described in Supplementary Fig. S1d-f, we obtained a reconstruction of TACAN with an overall resolution of 3.27 Å according to the 0.143 criterion of Fourier shell correlation (FSC) (Supplementary Fig. S1g). Local resolution analysis showed that most of the transmembrane region had a resolution within 3.0 and the intracellular region had a relatively lower resolution (Supplementary Fig. S2).

Our reconstruction revealed that TACAN formed a homodimer with the length and the axial height about 90 Å and 80 Å, respectively (Fig. 1a-d). The high quality of the density map allowed us to build an *ab initio* atomic model for the protein (Fig. 1e-g). Each monomer consists of a body (residues 123-249, 265-335), a spring (residues 101-122) and a blade (residues 7-100) domains. The body domain contains six transmembrane helices (TM1-6) that form a wedge-shaped channel with the wider end at the inner leaflet and the narrower end at the outer leaflet (Fig. 1e). A fenestration on one subunit connects the channel of the body domain with the membrane compartment (Fig. 1h). The blade domain has two helices (H1-H2) that contact with each other (Fig. 1e-f). The spring domain is composed of a short helix (H3) sandwiched by two loops with the helix running within and approximately parallel to the membrane (Fig. 1e-f). The transmembrane body and the cytoplasmic blade domains are linked together through connecting with the N-terminal and C-terminal loops of the spring, respectively (Fig. 1e-f).

**Fig. 1.**
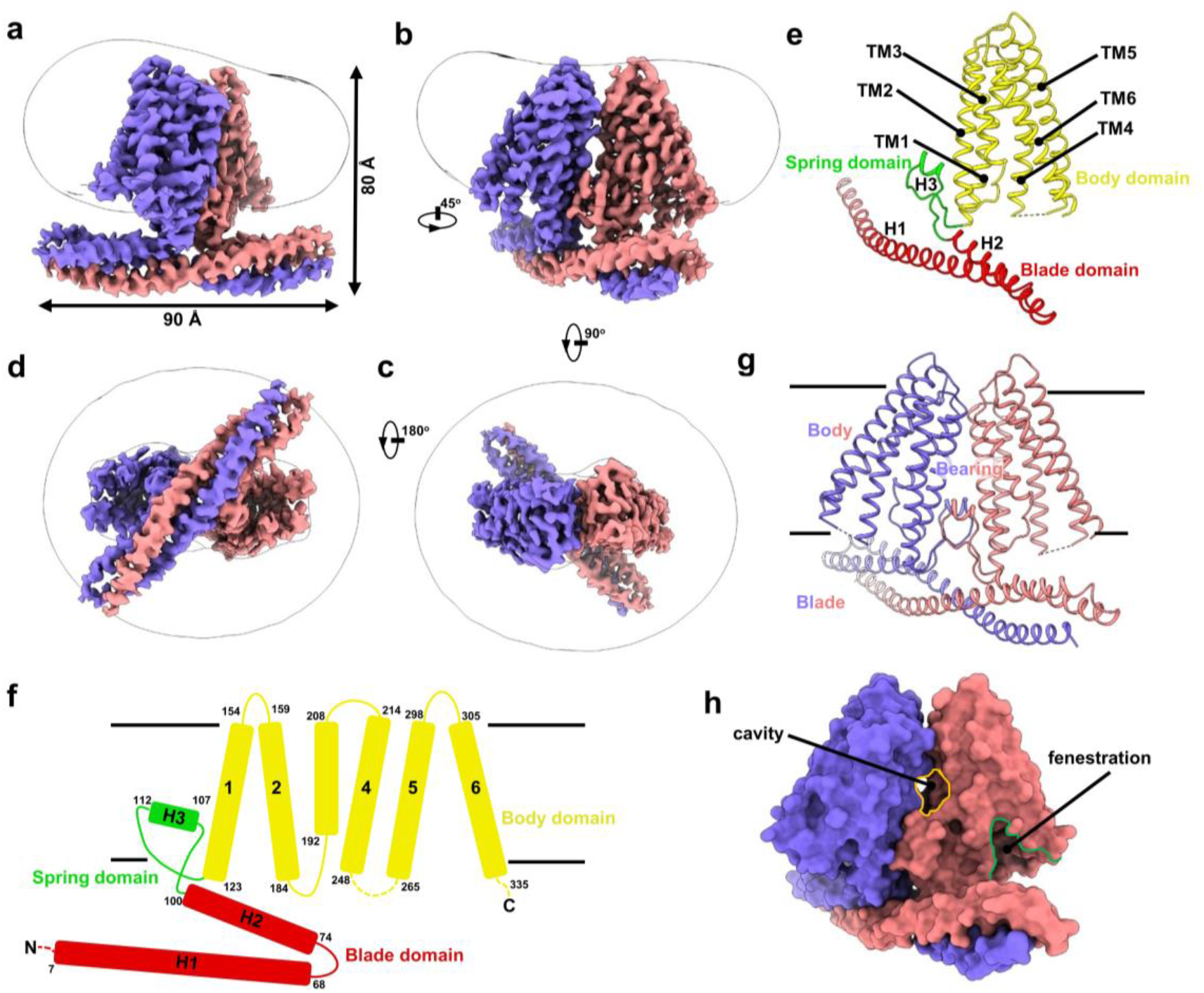
Overall structure of TACAN. **a-d** The side (**a, b**), top (**c**) and bottom (**d**) views of the density map. The two subunits of the TACAN homodimer are colored in purple and pink, respectively. **e** Atomic model of one subunit, colored by domain. **f** Topology and domain diagram of one subunit, colored by domain as in **e. g** A ribbon representation of TACAN structure. The two subunits are colored as the density maps in **a**-**d**. The two body domains form the vehicle body; the two spring domains form the vehicle bearing and the blade domains form the vehicle blade. **h** Surface representation of TACAN. The cavity residing between the two subunits is indicated by a yellow circle. The fenestration in one subunit is outlined in a green line.

### The interactions between the two monomers of TACAN

Each of the three domains from one subunit interact with its counterpart from another one (Fig. 2a). The two body domains contact each other only at the upper region proximal to the outer leaflet, thus sealed the extracellular side of the membrane (Fig. 2a). The body interaction is mediated mainly by a hydrophobic core that is formed by Val159, Phe166 and Val169 on TM2, Leu208 on TM3, Trp305 on TM6 and their symmetrical related residues labeled with an additional single quotation mark (‘) (Fig. 2b). The two spring domains insert into the large opening between the lower region of the body domains to seal the cytoplasmic side, but leave a cavity opens to the membrane (Fig. 2a). The spring domains are firmly held in place through hydrophobic interactions and hydrogen bonds with the body domains (Fig. 2c-d). While the configuration of the spring domain is maintained mainly by hydrogen bonds (Fig. 2e), the two spring domains have direct contacts through hydrophobic interaction mediated by a Leu and Val zipper (Fig. 2f). In addition, the two helices of one blade domain are twisted with the counterparts of another one to form a 4-helix bundle mainly through hydrophobic interactions and hydrogen bonds (Fig. 1g and 2g). The overall architecture of TACAN homodimer resembles a vehicle. The two body domains form the vehicle body; the two spring domains form the vehicle bearing and the blade domains form the vehicle blade (Fig. 1e-g).

**Fig. 2.**
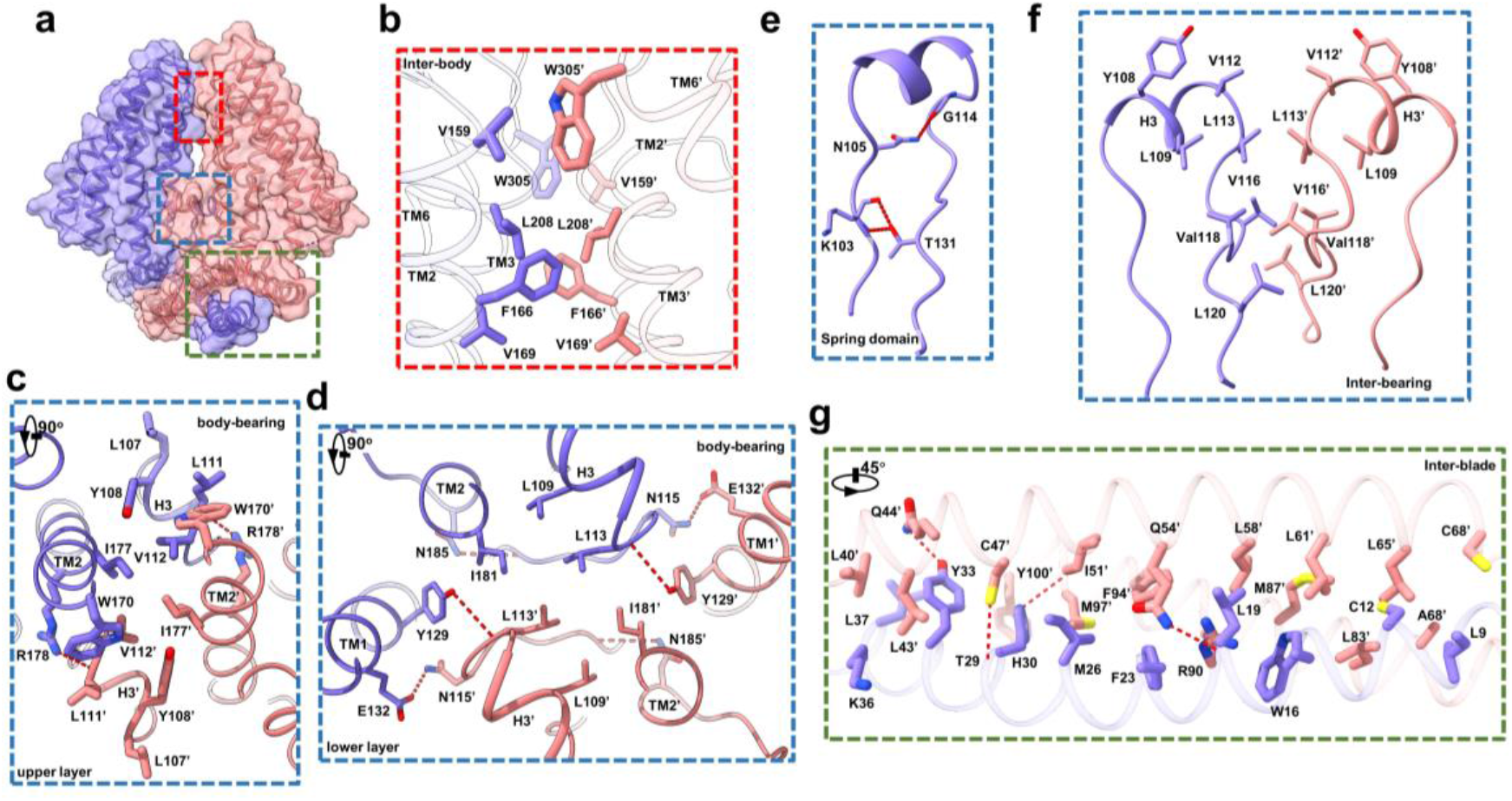
The interactions between the two monomers of TACAN. **a** Density map (transparent) and model (ribbon) of TACAN. The interactions between subunits are indicated by boxes. **b** Hydrophobic interaction between the two body domains of the red box region in **a. c**-**d** Hydrophobic interaction and hydrogen bonds (red dot lines) between the body and the bearing of the blue box region in **a**. The residues involved in hydrophobic interaction are indicated by sidechain showing only. **e** Hydrogen bonds that likely maintain the configuration of the spring domain. The hydrogen bonds are shown by red dot line. **f** Hydrophobic interaction between the spring domains of the blue box region in **a. g** Hydrophobic interaction and hydrogen bonds identified in the vehicle blade. For clarify, only one of the two symmetrical units of the blade is shown. Hydrogen bonds are indicated by dot line. The residues involved in hydrophobic interaction are indicated by sidechain showing only. The labels with and without single quotation mark (‘) are used to the distinguish the structural components or residues that are two-fold symmetrically arranged.

#### The closed channel

The entry pathway analysis showed that the body channel is constricted at the position formed by residues Asn165 on TM2, Met 207 on TM3 and Phe223 on TM4 (Fig. 3a-b). From the extracellular side, there are two pathways reaching to the constricted site (Fig. 3a), whereas from the intracellular side, there is only one gradually narrowing tunnel leading to the constricted site (Fig. 3a). Along this tunnel, several water molecules and one cation are tentatively assigned according to their densities and the surrounding polar residues (Fig. 3c), implying that the tunnel can communicate with the cytoplasm. Using Dali sever^13^, the body domain of TACAN is similar to the structure of the fatty acid elongase ELOVL7 (PDB 6Y7F)^14^ except the elongase has one more transmembrane helix of TM1 (Fig. 3d-f). However, TACAN doesn’t has the conserved motif HxxHH that is essential for the activity of fatty acid elongase (Fig.3f), which indicates that TACAN unlikely has the activity of the fatty acid elongase.

**Fig. 3.**
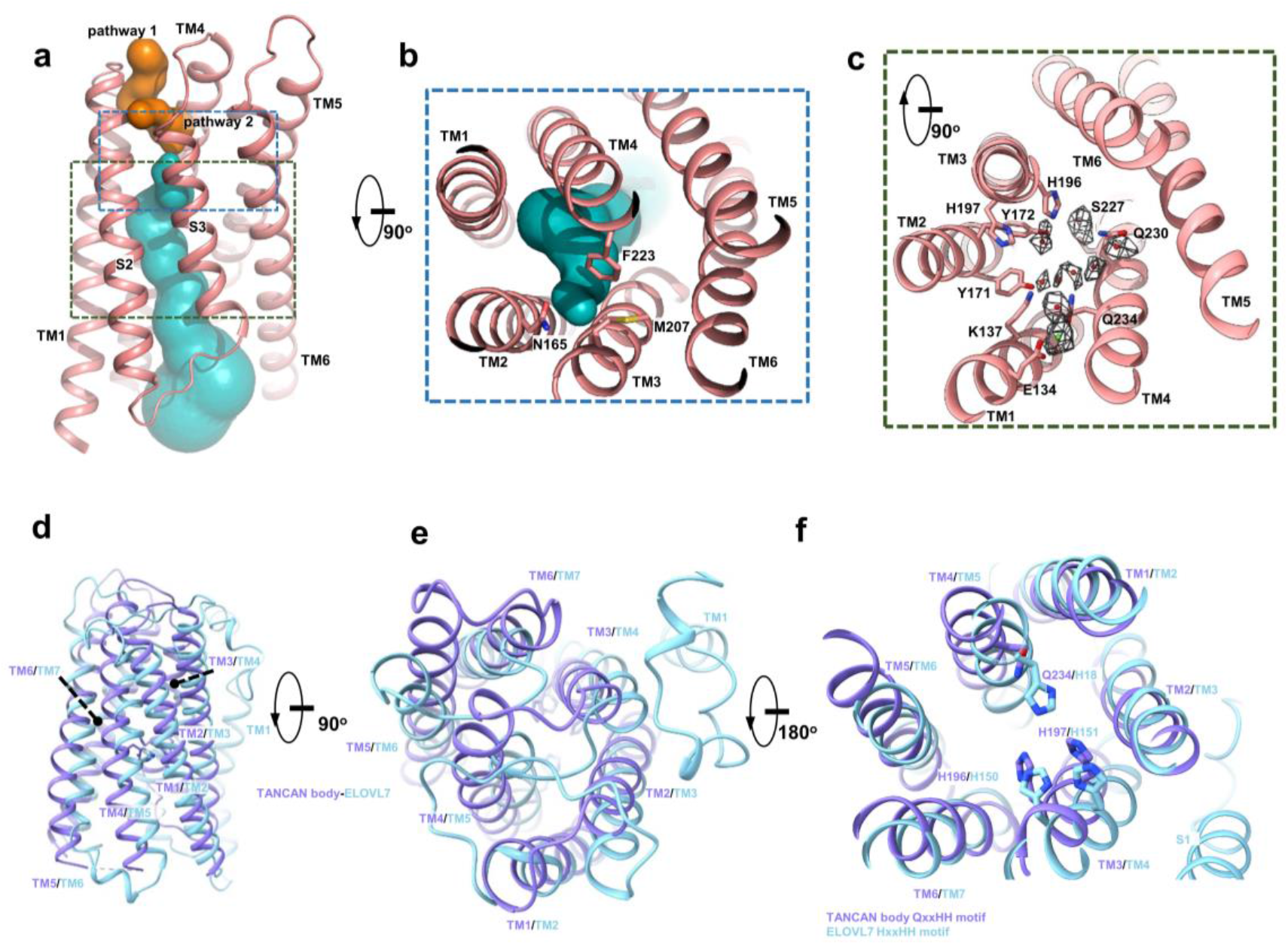
The channel of TACAN. **a** Side view of the channel, showing the channel is closed state and is interrupted at about one-third from the top. The upper part of the channel has two tunnels (orange). The lower tunnel is in cyan. The channel is calculated with MOLE 2.5^25^. **b** Top view of the channel, showing the channel is interrupted by Asn165, Met 207 and Phe 223. **C**, Top view of the lower tunnel, showing the possible H_2_O and cation densities. **d**-**f** Structural comparison between TANCAN body domain (purple) and fatty acid elongase ELOVL7 (cyan) viewed from side (**d**), top (**e**) and bottom (**f**). The key motif HXXHH for fatty acid elongase activity is missing in TACAN. The motif HXXHH of ELOVL7 and its corresponding residues in TACAN are indicated by sidechain showing (**f**).

#### TACAN is intensively associated with membrane lipids

Prominently, we found 11 strong lipids densities in our final reconstruction of TACAN, which are specifically associated with the helices of the body and spring domains (Fig. 4a-h). Lipid 1 interacts with two helices of TM4 and TM5 and is orientated parallel to the cell membrane. While the head of lipid 1 resides within a hydrophobic cavity formed by residues Phe281, Phe284, Trp285 and Phe288 on TM5, the tail of it extends to contact TM4 (Fig. 4a and 4f). Lipid 2, 3 and 4 form a linear cluster to contact with TM2, TM3, TM6 and TM1’ (Fig. 4a-b). In addition, a lipid core, formed by lipid 6, lipid 5 and its symmetrical molecule lipid 5’, resides within the cavity of the body (Fig. 4a-c). The densities of lipids are good enough for us to tentatively assign cholesterol for lipid 1, phosphatidylethanolamine (PE) for lipid 4, phosphatidic acids (PA) for lipid 5 and cardiolipin (CL) for lipid 2, lipid3 and lipid 6, respectively (Fig. 4d and Supplementary Fig. S2b). The phosphate heads of lipid 5 and lipid 5’ are attracted to the polar head of lipid 6 though hydrogen bonds (Fig. 4e). The lipid core symmetrically interacts with 4 helices of TM2, TM2’, H3 and H3’ mainly through hydrogen bonds (Fig. 4g). One of the two lipid acyl chain from lipid 5 extends upward into the hydrophobic zone formed by residues from TM1, TM2, TM1’ and TM2’ (Fig. 4h), whereas another lipid acyl chain stretches out from the cavity to contact the lipid cluster that is lying at the interface between the membrane and the protein (Fig. 4a-c and 4h), thus bridges the two linear clusters together.

**Fig. 4.**
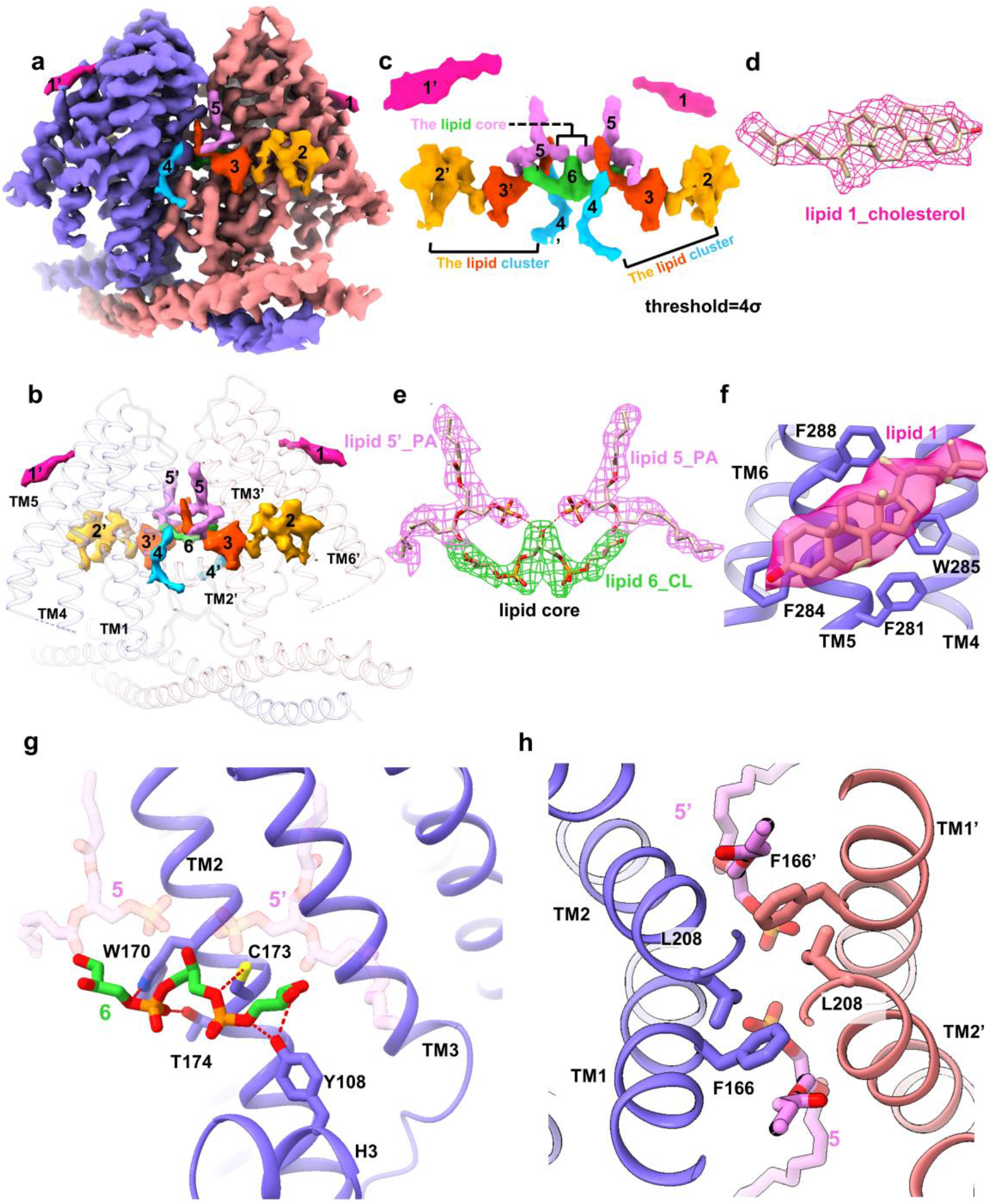
Membrane lipids associated with TACAN. **a** The density map of TACAN and the associated lipids. The two protein subunits are in purple and pink. The associated lipids are numerically labeled. The lipids that are two-fold symmetrically arranged are displayed with same colors but are distinguished by labels with and without single quotation mark (‘), respectively. **b** Atomic model (purple and pink ribbon) of TACAN and density maps of the membrane lipids (indicated by numeric label). **c** Density maps of the 11 lipids identified in the reconstruction of TACAN. **d** Atomic models (ribbon) and density maps (mesh) of membrane lipids, showing the lipids 1 fits well with the model of cholesterol. **e** Atomic models (ribbon) and density maps (mesh) of lipid core, showing the lipids 5 (or 5’) and 6 fit well with the models of phosphatidic acids (PA) and cardiolipin (CL), respectively. The lipid core is formed through hydrogen bonds between lipid 6 and lipids 5 (or 5’) (red dashed line). **f** Density map (transparent pink) and atomic model (stick) of cholesterol, which is accommodated by a hydrophobic pocket formed by residues from TM4 and TM5. **g-h** Interactions between the lipid core and TACAN. The lipid 6 is associated with TACAN mainly through hydrogen bonds (red dashed line) (**g**). For clarify, only one of two monomers is showed. Lipid 5 and 5’(stick) extends into the hydrophobic region of the TACAN body (ribbon) (**h)**. The labels with and without single quotation mark (‘) are used to the distinguish the structural components or residues that are two-fold symmetrically arranged.

## Discussion

Recently, one group has demonstrated that TACAN is a mechanosensitive ion channel and contributes to sensing the mechanical pain. Another group independently has shown that TACAN is significantly linked to the mechanical hyperalgesia produced by inflammation ^1, 2^. When we are preparing our manuscript, several groups presented the cryoEM structure of TACAN and showed that TACAN is not a mechanosensitive ion channel by electrophysiology^3, 4, 5^. Although there were these publications, our results are still significant in that the extensive and specific interactions between the membrane lipids and the membrane-embedded protein revealed in our study strongly imply that TACAN could sense the mechanical stimuli from the membrane lipids to perform biological functions. Additionality, it may provide an alternative mechanism how the TACAN responds to the mechanical stimulus. Actually, a previous study showed that the membrane lipids have the intrinsic feature leading to the possibility of soliton propagation in nerves^15^. Therefore, our results shall suggest that the TACAN may use membrane lipids to directly sense and transduce the mechanic stimulus, whereas the protein part of TACAN may function to modulate the action of lipids.

## Materials and Methods

### Protein expression and purification

The expression plasmid was reconstructed by cloning the cDNA of human TACAN into the pCAG vector with a carboxy-terminal Flag tag. HEK293F cells (Thermo Fisher Scientific) were maintained in Freestyle 293 medium (Thermo Fisher Scientific) at 37 °C in supplement with 5% CO_2_ and 80% humidity. When cell density reached 3.0 × 10^6^ cells/mL, the cells were transiently transfected with the expression plasmids using polyethylenimines (PEI, Polysciences). In details, about 1 mg expression plasmids were pre-mixed with 3 mg PEI in 50 mL fresh culture medium and incubated for 15 min. The mixture was then added to 1 L of cell culture. At 12 hours post transfection, 10 mM sodium butyrate was added to the culture medium and the culture temperature was shifted to 30 °C. Cells were collected 60 hours after transfection for the purification of TACAN.

Cell pellets were resuspended in the buffer containing 20 mM Tris-HCl pH 8.0, 150 mM NaCl and protease inhibitor cocktail (Bimake). The cells were lysed by high pressure homogenization and centrifuged at 8,000 g for 10 min at 4°C to remove the cell debris. The supernatant was further centrifuged at 100,000 g for 30 min at 4°C to enrich the membrane fraction. The membrane pellet was then solubilized using 20 mM Tris-HCl pH 8.0, 150 mM NaCl, 0.5% (w/v) lauryl maltose neopentylglycol (LMNG) (Anatrace) and 0.1% (w/v) cholesteryl hemisuccinate (CHS) (Anatrace) for 2 hours at 4 °C. Insolubilized material was removed by ultracentrifugation at 100,000 g for 30 min and the solubilized fraction was immobilized by batch binding to Flag affinity resins. The resins were then packed and washed with 20 column volumes of 20 mM Tris-HCl pH 8.0, 150 mM NaCl, 0.01% (w/v) LMNG and 0.002% (w/v) CHS. The target protein captured to the affinity resins was eluted using the washing buffer containing 200 ng/μl Flag peptide and concentrated using an 100kD Amicon Ultra Centrifugal Filter. The concentrate was subjected to size-exclusion chromatography (SEC) using a Superdex 6 Increase 10/300 column (GE Healthcare) equilibrated with buffer consisting of 20 mM Tris-HCl pH 8.0, 150 mM NaCl, 0.00015% LMNG and 0.00003% CHS to separate target protein from contaminants. Fractions from SEC were evaluated by sodium dodecyl sulfate polyacrylamide gel electrophoresis (SDS-PAGE) and those containing target protein were pooled and concentrated for cryo-EM experiments.

### Cryo-EM grid preparation and data acquisition

Peak fractions collected from SEC were concentrated to 18 mg/ml and then ultra-centrifuged at 40,000 g for 1 hour at 4 °C. Three microliters of the purified sample were applied onto a glow-discharged Quantifoil R1.2/1.3 300-mesh gold holey carbon grid. The grids were blotted for 3 s under 100% humidity at 4 °C and then plunge-frozen in liquid ethane cooled by liquid nitrogen using a Vitrobot Mark IV (Thermo Fisher Scientific). The frozen grids were carefully transferred and stored in liquid nitrogen.

A total of 23,964 movies were collected on a Titan Krios (FEI) equipped with a Gatan image filter and a K3 Summit detector (Gatan) operated at 300 kV accelerating voltage. SerialEM was used to automatically acquire micrographs at a nominal pixel size of 1.071 Å and defocus values ranging from −1.5 to −3 μm. Movies with 40 frames each were collected at a dose rate of 25 electrons per physical pixel per second for 3 s, resulting in a total dose of 70 e^-^/Å^2^ on the specimen.

### Image processing and 3D reconstruction

All 40 frames in each movie stack were aligned and dose weighted using MotionCor2^16^. Gctf was used for estimating the defocus values and astigmatism parameters of the contrast transfer function (CTF)^17^. A total of 20,097 micrographs were chosen from 23964 movies for further processing. About 5,000 particles were initially picked by ManualPick in RELION 3.1 from selected micrographs^18^. The particles were extracted on a binned dataset with a nominal pixel size of 2.142Å and were subjected to reference-free 2D classification. The reference-free 2D class averages selected from manually picking particles were used as templates for automated particle picking, which yielded a total of 15,974,407 particles. These particles were extracted on a 2 times binned dataset with a box size of 128 × 128 pixels and processed with 2D classifications. An initial reference model for 3D classifications was generated de novo from selected 2D particles using the stochastic gradient descent (SGD) algorithm. Finally, 198,882 particles from a 3D class showing good secondary structural features in the transmembrane domain were selected for further 3D refinement. After several iterations of CTF refinement and Bayesian polishing, we obtained a reconstruction with an overall resolution of 3.27 Å according to the Fourier shell correlation (FSC) equal to 0.143 criterion with a calibrated pixel size of 1.06 Å^19^. The local resolution was calculated by ResMap^20^.

### Model building and refinement

The atomic model of TACAN was built *de novo* using Coot ^21^. The real space refinement of the model was performed using PHENIX^22^. All of the structural figures were prepared using Chimera^23^, ChimeraX^24^ and PyMOL (https://pymol.org/2/). The final refinement statistics were provided in Table 1.

**Table 1.** Cryo-EM data collection, refinement and validation statistics.

| Data collection and processing |  |
| --- | --- |
| Magnification | 81,000 |
| Voltage (kV) | 30 |
| Electron exposure ($e^-/\text{\AA}^2$) | 70 |
| Defocus range ($\mu\text{m}$) | -2.5 |
| Pixel size ($\text{\AA}$) | 1.075 |
| Symmetry imposed | C2 |
| Initial particle images (no.) | 15,974,407 |
| Final particle images (no.) | 195,149 |
| Map resolution ($\text{\AA}$) | 3.2 |
| FSC threshold | 0.143 |
| Refinement |  |
| Model resolution ($\text{\AA}$) | 3.3 |
| FSC threshold | 0.5 |
| Map sharpening B factor ($\text{\AA}^2$) | -75 |
| Model Composition |  |
| Non-hydrogen atoms | 5260 |
| Protein residues | 626 |
| B factors ($\text{\AA}^2$) | |
| Protein | 61 |
| R.m.s.deviation |  |
| Bond length ($\text{\AA}$) | 0.004 |
| Bond angle ($^\circ$) | 0.615 |
| Validation |  |
| Molprobity score | 1.47 |
| Clashscore | 8.86 |
| Ramachandran plot |  |
| Favored (%) | 99.19 |
| Allowed (%) | 0.81 |
| Outliers (%) | 0.00 |

## Data availability

The density map has been deposited in the Electron Microscopy Bank under accession codes EMD-XXX. The atomic coordinate has been deposited in the Protein Data Bank under accession code XXX.

## Acknowledgements

The cryo-EM data were collected at Cryo-Electron Microscopy Research Center, Shanghai Institute of Material Medica. This work was partially supported by the Hundred Talents Program of Chinese Academy of Sciences (to X.Y.); Chinese Academy of Sciences grant (XDA12010317 to X.Y.); the Natural Science Foundation of Shanghai (18ZR1447700 to X.Y.); Shanghai Municipal Science and Technology Major Project (TZX022021007 to X.Y.); the National Natural Science Foundation of China (32000896 to Z.W.); China Postdoctoral Science Foundation (2020M681427 to Z.W.).

## Contributions

X.Y. and Z.W. conceived the research. X.Y. initiated and supervised the research. F.F., and Z.W. performed sample preparation. Q.W., Y.Z., F.F. and Z.W. collected the data. X.Y. and L.D. processed the data. L.D. and F.F. built the model. F.F. participated in data processing. R.G. and Z.W. analyzed the channel. X.Y. and Z.W. interpreted the data and wrote the manuscript.

