## Supplemental for "Cryo-electron microscopy revealed TACAN is extensively and specifically associated with membrane lipids"

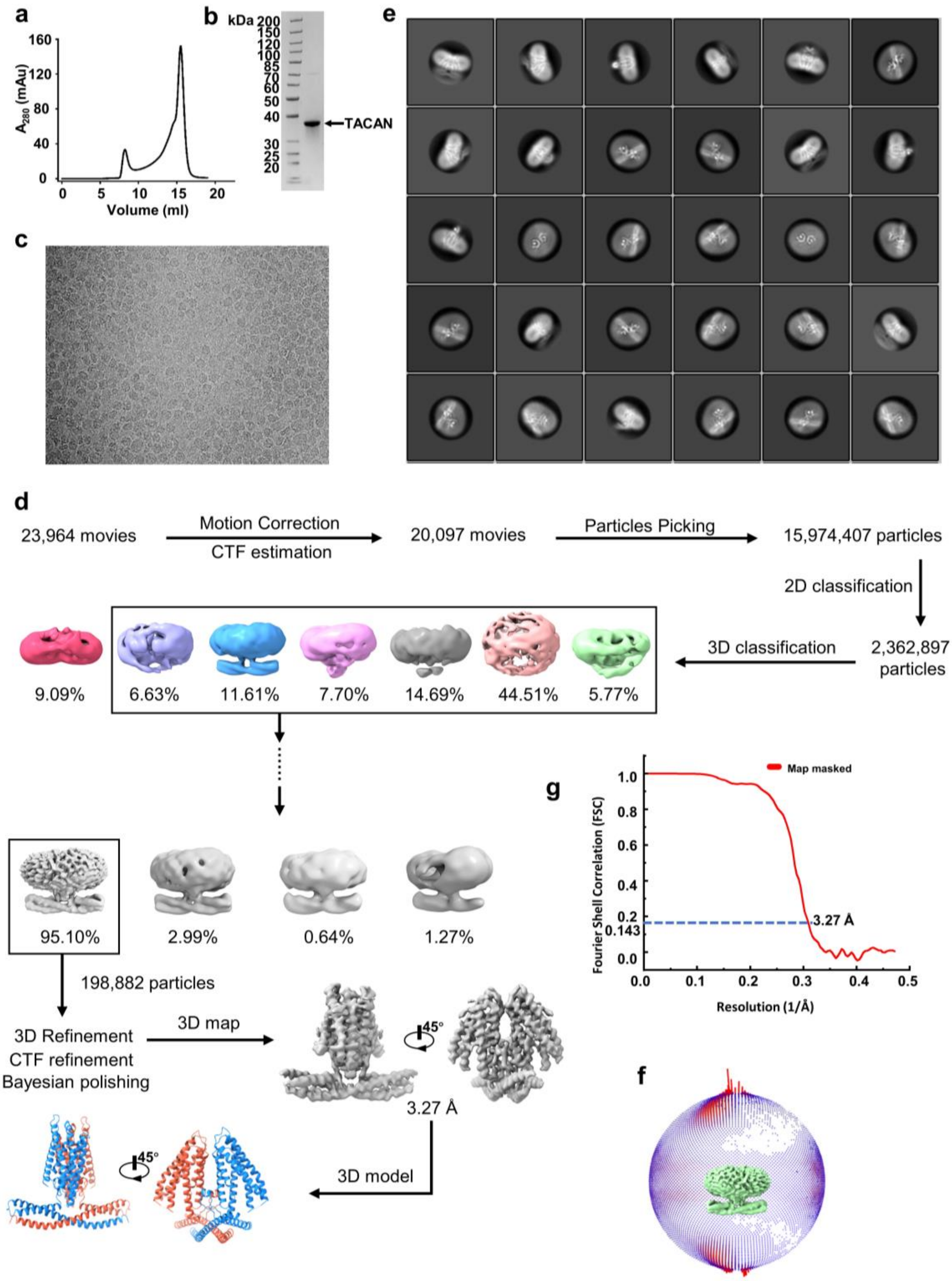


**Fig. S1 Sample preparation and image processing. a** Analytical size-exclusion chromatography purification of the expressed TACAN. **b** SDS-PAGE and Coomassie blue stain of the purified sample. **c** Representative cryo-EM micrograph. **d** Flow chart of image processing. **e** Representative 2D classes. **f** Angle distribution of particle images used for final reconstruction. **g** Gold-standard Fourier shell correlation (FSC) curves of the final 3D reconstruction.

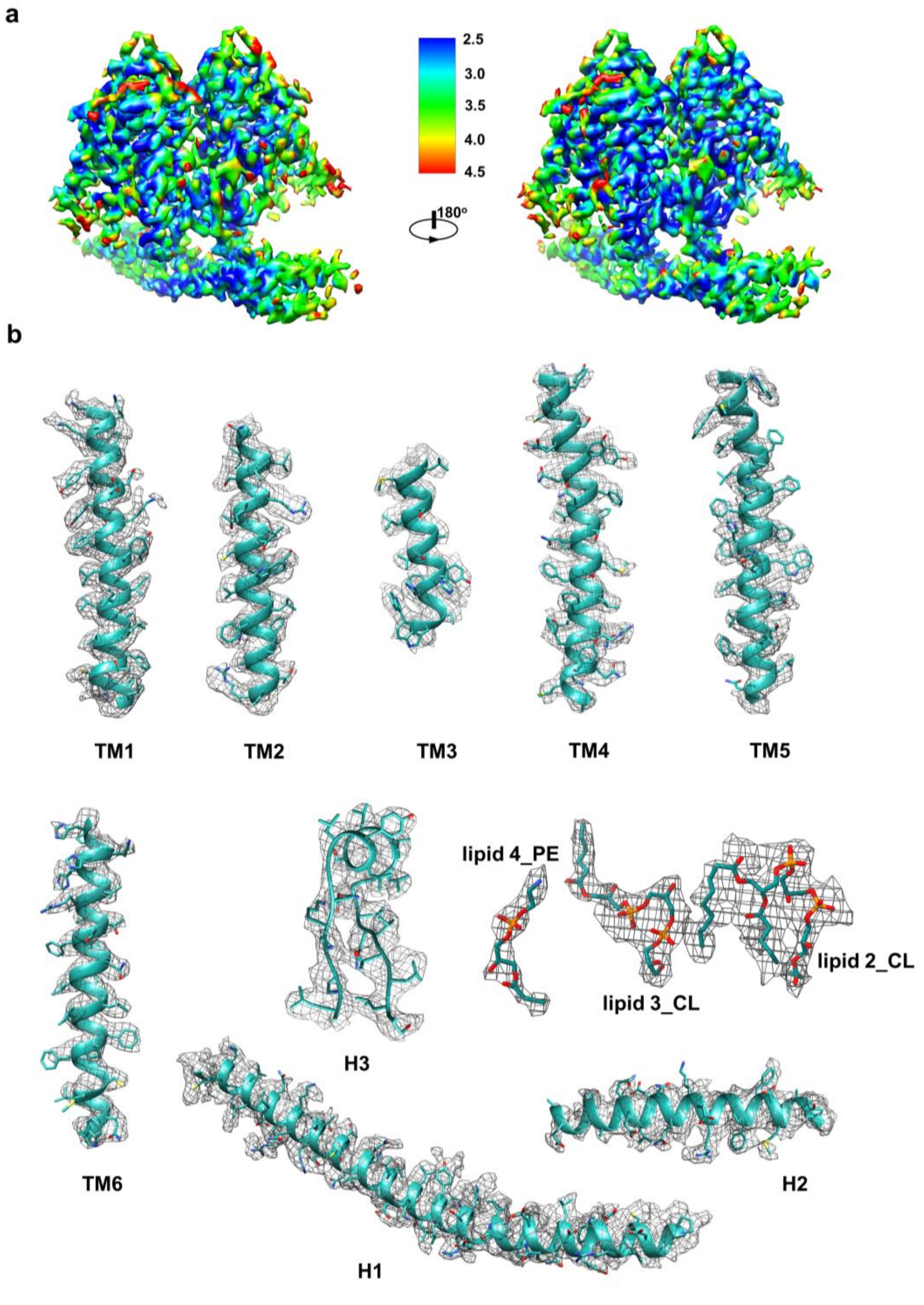


**Fig. S2 Local resolution and representative density maps of our reconstruction of TACAN. a** Local resolution distribution of the reconstruction estimated by ResMap. **b** Density maps (mesh) and models of helices (ribbon) and the lipids (stick).
